# Processing of grassland soil C-N compounds into soluble and volatile molecules is depth stratified and mediated by genomically novel bacteria and archaea

**DOI:** 10.1101/445817

**Authors:** Spencer Diamond, Peter Andeer, Zhou Li, Alex Crits-Christoph, David Burstein, Karthik Anantharaman, Katherine R. Lane, Brian C. Thomas, Chongle Pan, Trent Northen, Jillian F. Banfield

## Abstract

Soil microbial activity drives the carbon and nitrogen cycles and is an important determinant of atmospheric trace gas turnover, yet most soils are dominated by organisms with unknown metabolic capacities. Even Acidobacteria, among the most abundant bacteria in soil, remain poorly characterized, and functions across groups such as Verrucomicrobia, Gemmatimonadetes, Chloroflexi, Rokubacteria are understudied. Here, we resolved sixty metagenomic, and twenty proteomic datasets from a grassland soil ecosystem and recovered 793 near-complete microbial genomes from 18 phyla, representing around one third of all organisms detected. Importantly, this enabled extensive genomics-based metabolic predictions for these understudied communities. Acidobacteria from multiple previously unstudied classes have genomes that encode large enzyme complements for complex carbohydrate degradation. Alternatively, most organisms we detected encode carbohydrate esterases that strip readily accessible methyl and acetyl groups from polymers like pectin and xylan, forming methanol and acetate, the availability of which could explain high prevalences of C1 metabolism and acetate utilization in genomes. Organism abundances among samples collected at three soil depths and under natural and amended rainfall regimes indicate statistically higher associations of inorganic nitrogen metabolism and carbon degradation in deep and shallow soils, respectively. This partitioning decreased in samples under extended spring rainfall indicating long term climate alteration can affect both carbon and nitrogen cycling. Overall, by leveraging natural and experimental gradients with genome-resolved metabolic profiles, we link organisms lacking prior genomic characterization to specific roles in complex carbon, C1, nitrate, and ammonia transformations and constrain factors that impact their distributions in soil.

## Introduction

Grassland ecosystems cover 26% of all land area, store 34% of global terrestrial carbon, and comprise 80% of agriculturally productive land^1,2^. Therefore grasslands have a significant impact on global soil carbon storage, trace gas emissions, and economic productivity^1,2^. Thus, it is critical that we identify organism capacities for carbon and nitrogen turnover, as microorganisms ultimately determine how grassland soils cycle carbon and nitrogen, and emit or absorb trace gases^3,4^.

One of the biggest challenges in studying the metabolism of soil microbial communities is that most of the organisms have only been detected using 16S rRNA surveys^5,6^. While multiple studies have been undertaken to link amplified metabolic genes or 16S rRNA gene abundances with soil trace gas fluxes or environmental conditions^7-11^, the large number of soil-associated bacteria that are not represented by genomes precludes meaningful predictions of relationships between organism types and their biogeochemical functions.

The metabolic capacities of soil-associated microorganisms can be investigated if genomes can be reconstructed from soil samples^12-14^. However, this is notoriously difficult, as many soils contain a vast diversity of organisms^15^. To date, few soil datasets have been even partially genomically resolved^13,16^, but recently it was shown that broad genomic resolution and community metabolic functions could be deduced in large scale metagenomic studies targeting permafrost^12^.

Here we applied deep metagenomic sequencing and metaproteomic analyses to sub-root zone samples from a grassland soil ecosystem from a Mediterranean climate. Mediterranean grassland soils are of particular interest as they can discharge a CO_2_ pulse following the first Fall rainfall equal to the annual CO_2_ output from many other ecosystem types^17,18^. The soils in this study were manipulated to test a long-term rainfall alteration climate change scenario^19^. Despite the presence of thousands of species at low abundance levels and strain heterogeneity, we successfully reconstructed non-redundant draft-quality genomes that account for the majority of organisms by abundance. Overall our data reveal important carbon and nitrogen turnover functions in understudied microbial groups, show a stark metabolic and phylogenetic stratification across soil depths, and support long term climate change as a factor that can significantly alter the carbon and nitrogen turnover capacity of soil microbial communities.

## Results

### Soil sampling and assembly

We collected 60 soil samples from 10-20 cm (just below the root zone), 20-30 cm, and 30-40 cm from a grassland meadow within the Angelo Coastal Range Reserve in Northern California (Supplementary Fig. 1). Three of the six sampling sites had been subjected to over 13 years of rainfall amendment to simulate a predicted climate change scenario for northern California^19^. In total, we generated 1.2 Tb of raw read data which assembled into 67 Gbp of contiguous sequence. Of this, 47 Gbp (70.2%) of the assembled sequences were >1 kb in length. Of all reads sequenced, 36.4% mapped back assemblies on average, and for some samples this mapping was as high as 64.7%. A total of 59 million genes were predicted on all contigs >1 kb (Supplementary Table 1).

### A species richness census reveals extensive sampling of soil microbial diversity

Although our approach overall is genome-centric, many organisms were at too low abundance to be represented by draft genomes. Thus, we used ribosomal protein S3 (rpS3) to conduct a census of the microbial diversity found at the site and to quantify global organism abundances^20^. Across our 60 metagenomic assemblies we identified 10,158 rpS3 sequences (169±93 per sample), which were grouped into 3325 non-redundant clusters (99% amino acid identity) that approximate species groups (SGs) (Supplementary Table 2 and Supplementary Data 1-2).

Our SG survey indicated that these soils are heterogeneous with a high prevalence of relatively low abundant organisms, as 2120 (63.7%) SGs were only assembled from one of our 60 metagenomic samples. However, by cross mapping reads from all 60 samples back to our representative SG sequences, we could detect and track the presence of a SG in a sample even when it was below the ~2x coverage threshold required for assembly (Supplementary Fig. 2a-b). We found that the 2120 sequences reconstructed in only one sample could be confidently detected on average in 31±18 samples at low abundance (Supplementary Table 2 and Supplementary Fig. 2b).

The iChao2 metric^21^ and a permuted collectors curve were used to estimate species richness and assess the impact of possible further sampling on novel SG recovery, respectively (Supplementary Fig. 2c-e). The iChao2 metric estimated total species richness at 9183 organisms (95% CI: 8641 - 9780 organisms). Thus, our SGs represent 34% - 38% of the total organisms present at this site by number. Our collectors curve indicated that we did not saturate SG recovery, however the slope of the recovery curve indicates that we have saturated high level recovery and further sampling would only yield around 30 new SGs per sample (Supplementary Fig. 2d-e).

Using our rpS3 sequences as phylogenetic markers we initially classified all of the organisms detected at the phylum and class level. We detected 26 distinct Phylum-level lineages, and the topology of the rpS3 tree suggested that most phyla are represented by just a few class level groups with high degrees of genus, species, and strain heterogeneity. We also found that the abundances of closely related organisms could be highly variable, differing in abundance >10-fold (Supplementary Fig. 3 and Supplementary Data 3).

In agreement with many previous soil surveys^5,6^, we found that Verrucomicobia and Acidobacteria were the most abundant lineages across our site (Fig. 1a). Some individual organisms had high relative abundance across the site, but the vast majority of organisms are present at low abundance. Coverage was disproportionately concentrated in a smaller subset of SGs, and approximately 13% (443) of the detected organisms accounted for 50% of the global read coverage (Fig. 1b). Some organisms, such as specific Nitrospirae and Euryarchaeota, were highly abundant although their phylum was not well represented overall (Fig. 1a). Thus, while some phyla do not collectively account for a highly fraction of the soil microbiomes, individual organisms belonging to these phyla may be highly abundant.

**Figure 1.**
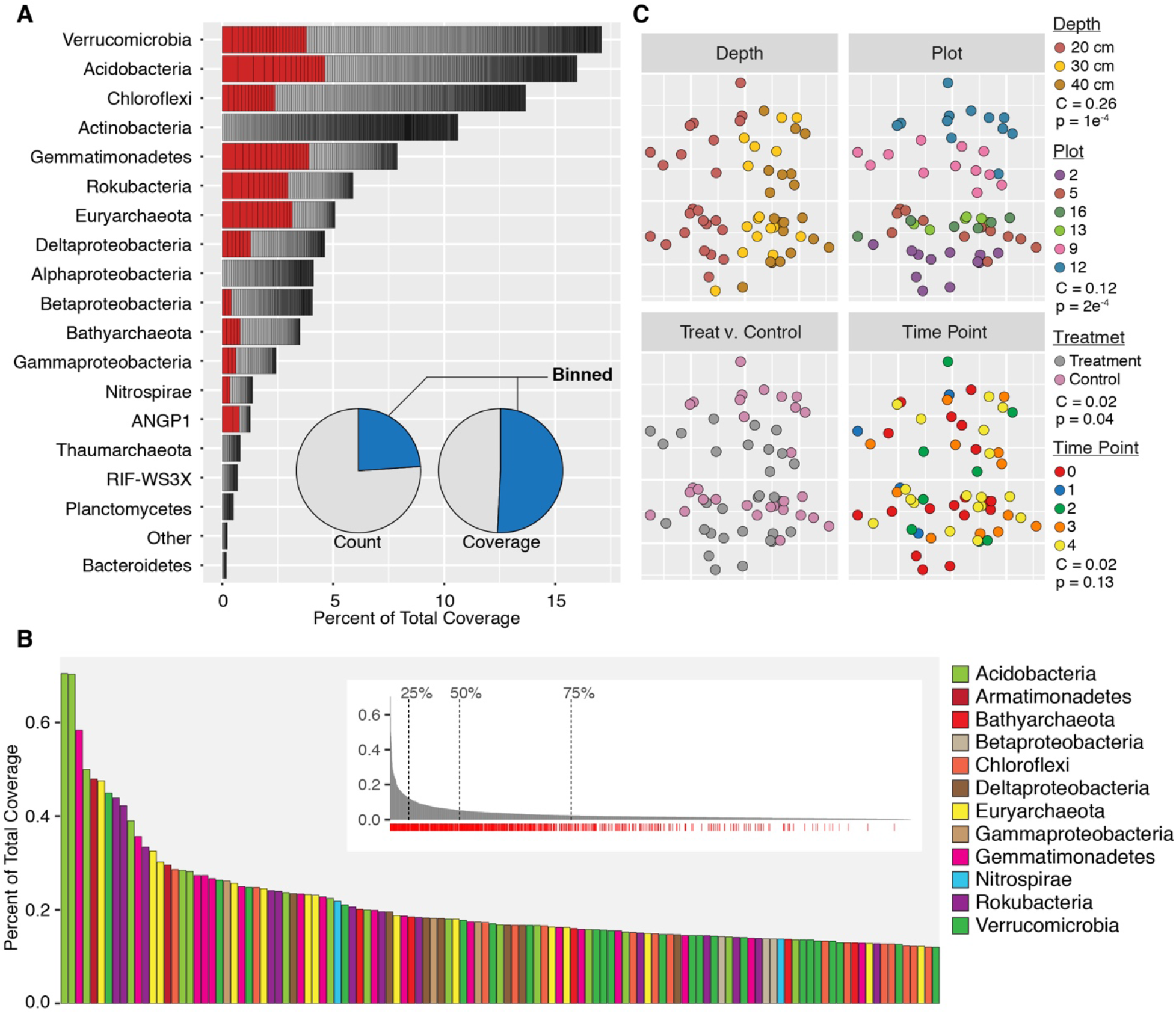
rpS3 species group abundance, influence of variables, and abundance metrics. (**A**) Percent of total coverage of all SGs ranked by total phylum coverage. “Other” includes phyla with < 5 SGs. Organisms in red are in the top 25% of organisms by coverage. The inset pie chart shows the breakdown of SGs associated with genome bins (blue) based on count and coverage of SGs. (**B**) NMDS plot (stress = 0.055) of SG UniFrac distances. The ordination is replicated and overlaid with the four data types collected across our 60 samples. Variable importance (C) and significance (p) calculated by MRPP are displayed in the legend. (**C**) Top 25% of SGs ranked by total coverage across all samples. The inset shows the full rank abundance curve and shows the positions where 25%, 50%, and 75% of the total dataset coverage are reached. Red ticks under the plot indicate SGs with bins.

### Spatial variation and treatment, but not time of sampling, contribute significant variance to organism abundance

To visualize the influence of depth, sampling location, sampling date, and rainfall amendment on the abundance of SGs, we applied non-metric multidimensional scaling (NMDS) ordination to the weighted Unifrac distance matrix of SG coverage (Fig. 1c, Supplementary Table 3-4, Supplementary Data 4). Subsequently we used the Multi-Response Permutation Procedure (MRPP) to test the significance and strength of each variable’s influence. The results indicate that sampling depth, sampling location, and rainfall amendment had significant effects on overall organism abundance and composition across samples (Fig. 1c). Sampling depth was the most influential factor (C = 0.26; p = 1e^-4^), followed by sampling location (C = 0.12; p = 2e^-4^), and rainfall amendment (C = 0.02; p = 0.04). While rainfall amendment showed a consistent effect, its effect occurs relative to sampling location (Fig. 1c). A large sample number was critical in observing this relationship, and was important for isolating the weaker effect caused by rainfall extension. Alternatively, we found that the date a sample was collected did not significantly influence overall SG variability, despite samples being collected over a 31-day period covering the transition from the dry to rainy season (Fig. 1c and Supplementary Fig. 1).

### A Hybrid binning method resolved genomes from previously unsequenced lineages

Genomes reconstructed from each sample were used to link metabolic functions to specific organisms (Methods). We recovered 10,463 genomic bins with an average of 174±87 binned genomes per sample. After clustering bins based on the SGs assigned to their rpS3 gene, and filtering for estimated completeness > 70% and contamination < 10%, we recovered 793 unique microbial genomes (Supplementary Table 5).

Our reconstructed genomes represent 24% of SGs by number, however these genomes represent more than half (53%) of the SGs by total coverage (Fig. 1a). 204 genomes were from organisms in the lowest quartile of global abundance (Fig. 1b). Importantly, we recovered 115 high quality genomes (> 95% estimated completeness) across 15 of the 26 microbial phyla detected at the site (Supplementary Table 5). No genomes were recovered for SGs from the Aenigmarchaeota, Elusimicrobia, Saccharibacteria, Microgenomates, and Parcubacteria groups, all of which had ≤ 3 detected SGs and occurred at very low abundances.

A more detailed phylogenetic analysis using both a concatenated set of 15 ribosomal proteins (rp15) and 16S rRNA sequences indicated that we have significantly expanded the genomic coverage across a number of poorly sequenced soil lineages (Fig. 2, Supplementary Fig. 4-5, Supplementary Table 5-6, and Supplementary Data 5-8). Many genomes from unsequenced lineages were highly abundant organisms at our site. In particular, we recovered 145 near complete Acidobacterial genomes from 15 class-level lineages, four of which have no previously sequenced representative (Gp18, Gp5, Gp11, and Gp2) (Fig. 2 and Supplementary Fig. 4-5). We also found phylogenetic overlap between our Acidobacterial genomes and previously recovered but unclassified Acidobacterial genomes from a subsurface aquifer sediment in Rifle, Colorado^22^. By including genomes from both the Rifle and Angelo sites in our phylogenetic tree we were able to assign 17 genomes to Acidobacterial Classes Gp7, Gp22, and Gp17 for which there was no previous class-level genomic information (Supplementary Fig. 5).

**Figure 2.**
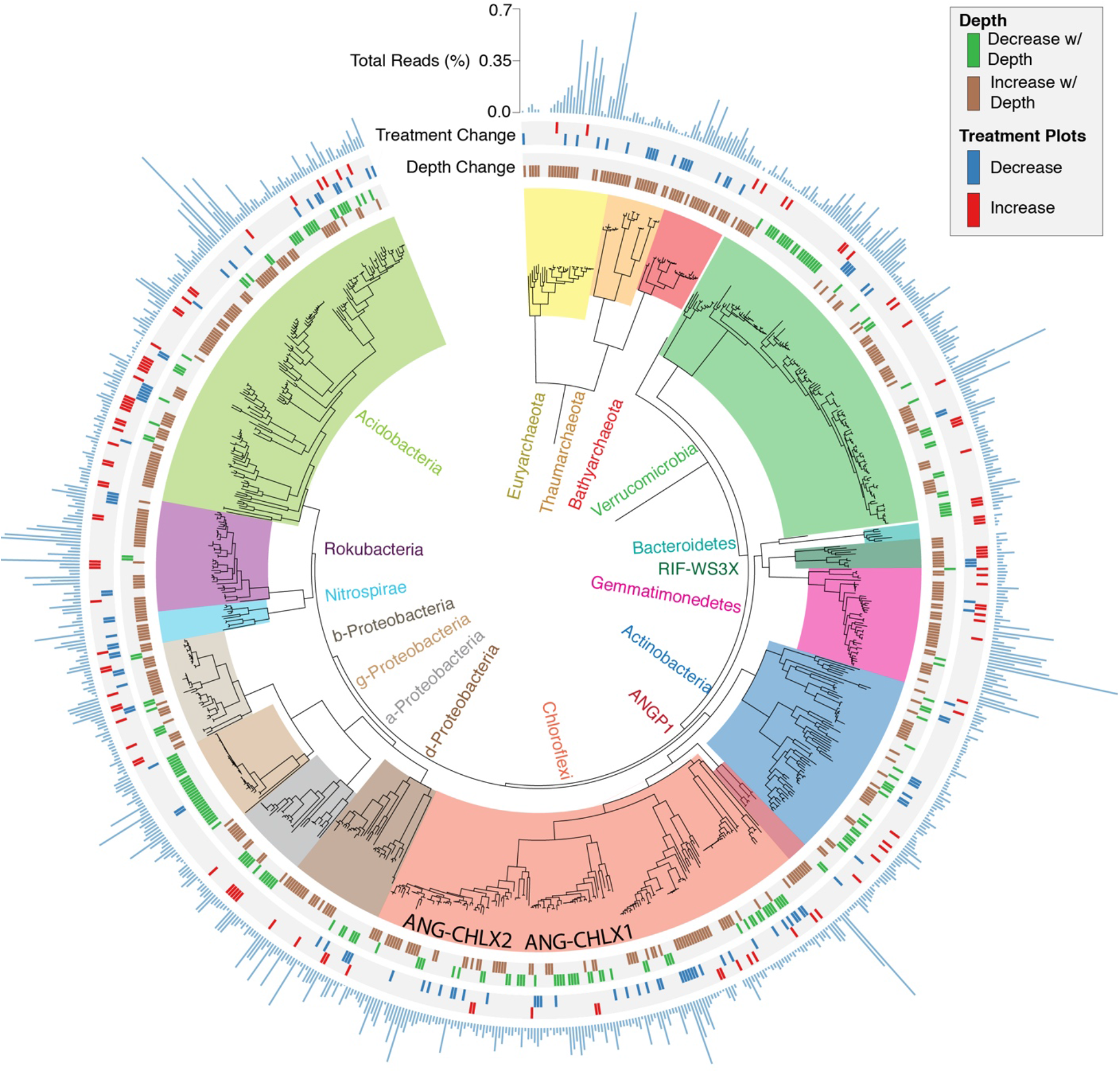
Maximum likelihood tree of all near complete genomes. Phylogenetic tree constructed with a concatenated alignment of 15 co-located ribosomal proteins. Tree includes 722 Bacterial and 71 Archaeal genomes. The two novel Chloroflexi classes are named. Concentric rings moving outward from the tree indicate if a genome’s associated SG was found to increase or decrease with depth and increase or decrease with treatment at 10-20 cm and 30-40 cm, respectively. The concentric bar plot indicates relative abundance (Methods). For the complete ribosomal protein tree, see Supplementary Fig. 4 and Supplementary Data 5.

The majority of our Chloroflexi genomes came from four unsequenced or poorly sequenced approximately class-level lineages. Nine genomes affiliate with a group referred to as CHLX from Rifle aquifer sediment^22^, and 32 genomes phylogenetically place with a second lineage that includes one genome from Rifle sediment and one from arctic soil^13^. We also recovered 96 genomes from two class-level lineages within Chloroflexi with no previously sequenced representatives, hereafter referred to as ANG-CHLX1 and ANG-CHLX2 (Fig. 2, Supplementary Fig. 4). The ANG-CHLX1 and ANG-CHLX2 clades form a strongly supported group basal to RIF-CHLX genomes and all known Chloroflexi lineages.

### The soil proteome indicates a high prevalence of C1, pentose sugar, and small molecule metabolism

We used shotgun proteomic data from 20 samples to provide insight into abundant functions in situ (Methods), and to guide or metabolic analysis of the reconstructed genomes. Overall, we identified 55,665 proteins with at least one uniquely mapped peptide that was detected with high mass accuracy. To enable comparison across samples, proteins from each sample were assigned and clustered into functional orthology groups. In total, 63% of the proteins identified could be assigned to one of 1,114 orthology groups (Supplementary Tables 7-8 and Supplementary Data 9). Additionally, we observed that the top 50 functions account for 62% of annotated proteins, suggesting that a small set of metabolic enzymes are particularly important in this system (Supplementary Fig. 6).

The most abundant proteins identified were associated with ABC transport of sugars and amino acids, the processing of pentose sugars, and the degradation of small C1 and nitrogen containing compounds (Supplementary Fig. 6). While we expected to find numerous transporters, pentose processing metabolic genes, and had previously detected a high abundance of methanol dehydrogenases at this site^14^, we were surprised by the high abundance of CoxL-like carbon monoxide dehydrogenases and enzymes like formamidase dedicated to small nitrogenous compound degradation (Supplementary Fig. 6). As our proteomics results indicated the importance of low molecular weight compound metabolism, we specifically targeted these enzyme types, as well as general C1 metabolic pathways, in our subsequent analysis of genome metabolic potential.

### The analysis support the importance of metabolism of small molecule in soil and identify nitrogen cycling processes in unexpected organisms

The dbCAN and KEGG databases were used to profile soil genomes^23,24^, as well as specific pathways selected based on the proteomics data (See Methods, Supplementary Tables 9-12, and Supplementary Data 10-11). Additionally, we note that methanol dehydrogenase (xoxF), CO dehydrogenase (coxL), and nitrite reductase (nirK) enzymes were classified both by homology and by using phylogenetically reconstructed gene sets (Supplementary Figs. 7-9, Supplementary Table 13, and Supplementary Data 12-14).

187 genomes encode a methanol dehydrogenase, and all methanol dehydrogenases identified in our genomes were of the XoxF-type (Fig. 3a, Supplementary Fig. 7). Consistent with a previous study at this site^14^, these enzymes were highly prevalent in Gemmatimonadetes and Rokubacteria, but also were detected in Gp1, Gp5, and Gp6 Acidobacterial genomes and 4 phyla of Proteobacteria (Fig. 3a). 90 genomes encode formamidase (amiF), including 26 Chloroflexi and 30 Rokubacteria (Fig. 3a). Formamidase contributes to both formate and ammonia pools via the breakdown of formamide. Formamide in uncontaminated soils might be derived from breakdown of amino acids via pathways with cyanide intermediates^24^. Using coxL as a marker for coxLMS type CO-dehydrogenases^9^ we detected 1889 coxL homologues encoded in 466 genomes. However, only type-I of the enzyme is known to be strongly active on CO^9,25^. Phylogenetic classification indicated that the vast majority of these proteins represented subtype-II and other novel groups of CoxL-like dehydrogenases (Supplementary Fig. 8). However, of the 100 genomes encoding type-I coxL genes, 59 were from the phylum Chloroflexi, with the majority being part of the ANG-CHLX1 and ANG-CHLX2 clades sequenced for the first time in this study (Fig. 3a).

**Figure 3.**
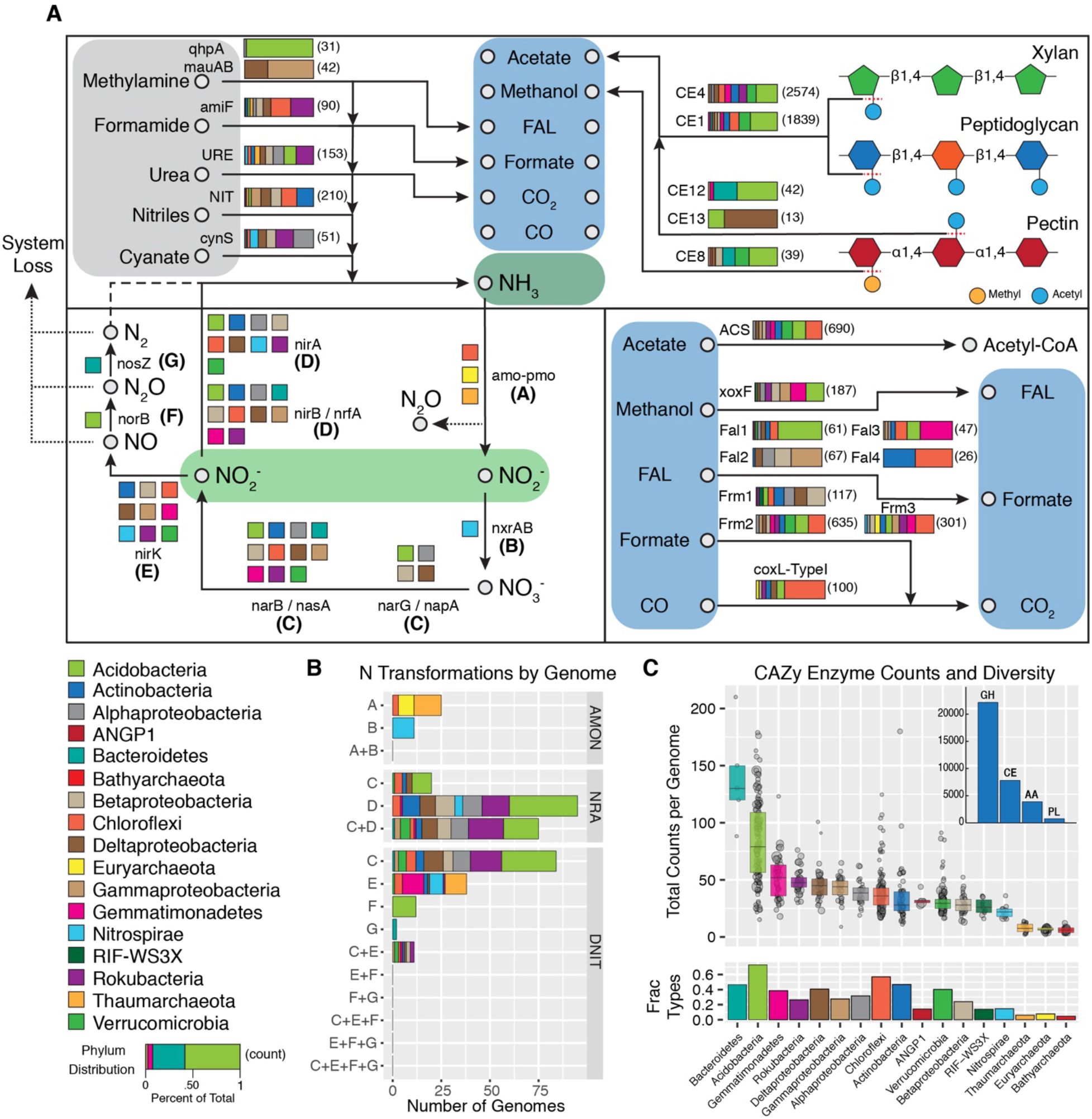
Predicted carbon and nitrogen metabolic transformations. (**A**) Top panel: predicted phylum level genomic capacity for breakdown of small carbon- and nitrogen-containing compounds. Bar plots indicate the fraction of genomes within a phylum encoding each function. Numbers adjacent indicate the total genes detected in 793 analyzed genomes. The derivation of small molecules via liberation of methyl and acetyl groups from complex polymers is illustrated. (**B**) Counts of genomes encoding capacities for individual or multiple nitrogen transformation steps. AMON = Ammonia Oxidation to Nitrate; NRA = Nitrate Reduction to Ammonia; DNIT = Denitrification. (**C**) Counts of carbohydrate active (CAZy) enzymes across genomes in each phylum. Points indicate the total counts in individual genomes and point sizes reflect genome relative abundance across all samples. The inset bar plot shows the total number of CAZy enzymes across all genomes belonging to each CAZy class (GH = glycosyl hydrolase; CE = carbohydrate esterase; AA = auxiliary activity; PL = polysacharide lyase). The fraction of all 246 possible CAZy enzymes types that were identified across the phylum is shown below. Also see Supplementary Tables 10-13.

Bacteroidetes and Acidobacteria genomes encode the largest number of CAZy enzymes, but Acidobacteria far exceed Bacteroidetes in terms of both total genomes detected (152 vs 5) and relative abundance (16% vs 0.2% across all communities) (Fig. 2, 3c, Supplementary Fig. 4). Acidobacteria also have the highest diversity of CAZy enzyme types, with 73% of CAZy families identified in this study being detected in at least one member of this phylum (Fig. 3c). A previous analysis of Acidobacterial genomes indicated that organisms from classes Gp1 and Gp3 contain large numbers of genes for complex carbohydrate degradation^26^. In this study, we identified 9 Acidobacterial classes containing genomes that encode >100 CAZy enzymes, including the previously unsequenced classes Gp2, Gp11, and Gp18 (Supplementary Fig. 10). This significantly expands the metabolic potential for complex carbohydrate turnover across the Acidobacteria phylum.

Across CAZy enzymes we noted a particularly high proportion of carbohydrate esterases (22%; Fig. 3a, 3c, and Supplementary Fig. 10). Types CE1 and CE4 account for 56% of all carbohydrate esterases identified, and liberate acetate from a broad spectrum of complex plant and microbial polymers^27,28^. Of the 793 genomes analyzed 93% contain either CE1 or CE4, and 81% of genomes contain one of the CEs as well as an encoded acetyl-coA synthetase to incorporate liberated acetate (Supplementary Fig. 10)

In a targeted analysis assessing genomic capacity to mediate inorganic nitrogen transformations we found most organisms encode genes for only a single transformation reaction, and that nitrite is the most common reaction substrate (Fig. 3a, 3b, and Supplementary Table 11). We did not detect any organism with the potential for complete denitrification, or complete nitrification via ammonia oxidation (Fig. 3b). Also, we found only two genomes classified as Bacteroidetes encoding the enzyme nosZ, which may indicate limited N_2_O turnover capacity in this system (Fig. 3b).

The phylogenetic affiliation of organisms encoding nitrogen transformation functions were in some cases surprising. Of the 49 genomes encoding nirK, 24% were from Gemmatimonadetes, a genomically undersampled phylum that is not normally linked to nitrite conversion to nitric oxide (Fig. 3b). Many Gemmatimonadetes with nirK were found in moderately high abundance (Supplementary Table 10). The gene norB, which converts nitric oxide (NO) to N_2_O, was exclusively found within genomes of Acidobacteria (Fig. 3b). While five Acidobacterial lineages had been reported to encode norB^29^, we additionally detected these genes in Acidobacteria from Gp4, Gp5, and Gp13 suggesting a widespread capacity for nitric oxide reduction across the Acidobacterial phylum (Supplementary Tables 5 and 10).

### Organisms are phylogenetically and functionally stratified by depth

391 genomes significantly increased and 179 decreased in abundance with increasing soil depth. Thus, the majority of assembled genomes (72%) exhibit abundance patterns stratified by depth (Fig. 2 and Supplementary Table 5). All Archaeal lineages as well as Rokubacteria, Gemmatimonadetes, Nitrospirae, and the candidate phylum RIF-WS3X were preferentially enriched in deeper samples, whereas Gammaproteobacteria were enriched at shallower depth (Fig. 4a, and Supplementary Table 14).

**Figure 4.**
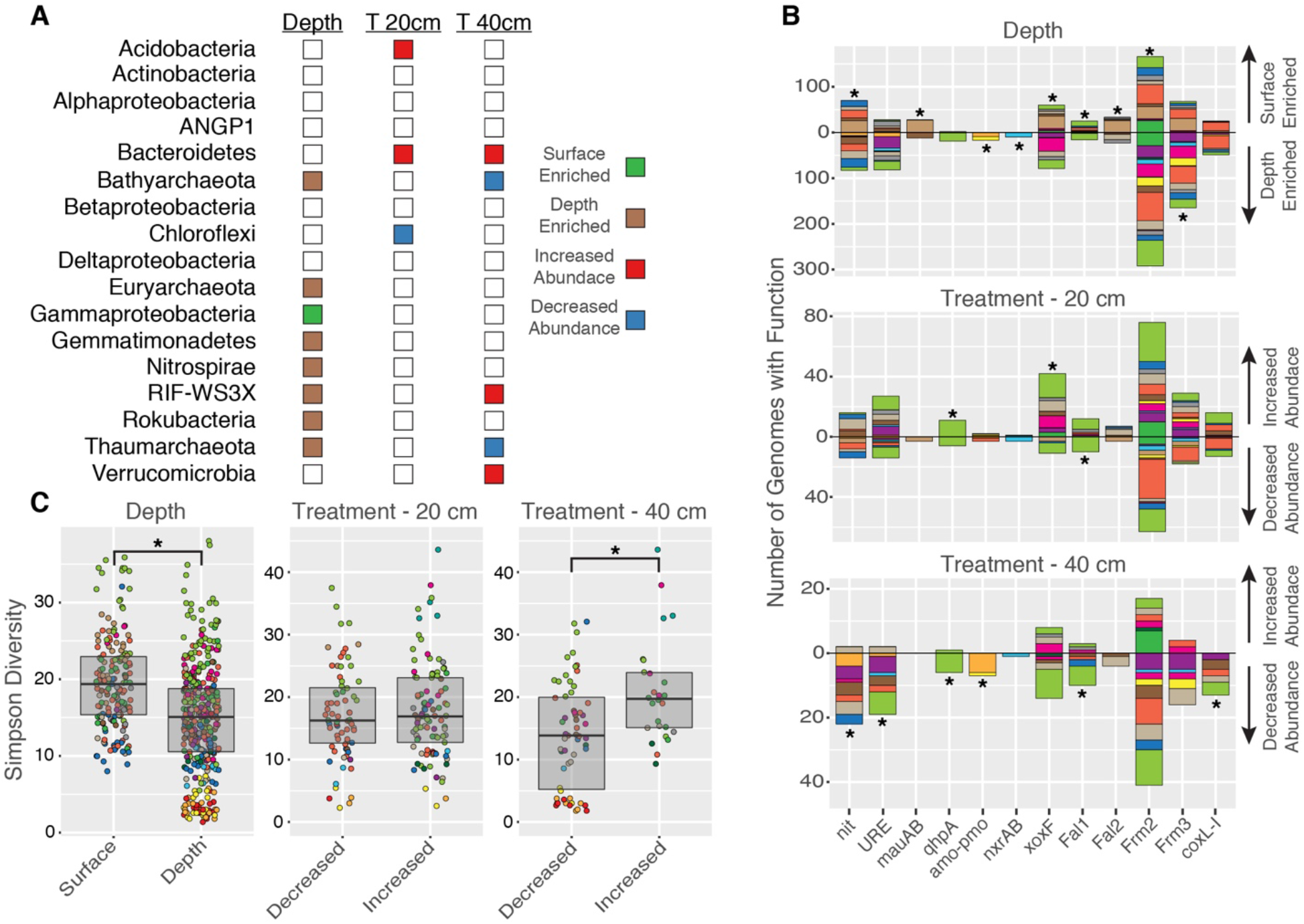
Enrichment of phyla and metabolic functions across depth and treatment. (**A**) Colored boxes indicate a significant enrichment (p <0.05; hypergeometric test); empty boxes indicate no effect. (**B**) The number of genomes encoding targeted carbon and nitrogen processing functions found to be significantly enriched in at least one comparison. Colors indicate phyla (see Fig. 3). Black stars indicate a significant enrichment of the function (p <0.05; hypergeometric test). Nitrilase (NIT); urease (URE); methylamine dehydrogenase (mauAB); quinohemoprotein amine dehydrogenase (qhpA); ammonia monooxygenase - particulate monooxygenase (amo-pmo); nitrite oxioreductase (nxrAB); methanol dehydrogenase (xoxF); formaldehyde oxidation - direct (Fal1); formaldehyde oxidation - glutathione pathway (Fal2); formate oxidation - multisubunit (Frm2); formate oxidation - THF mediated (Frm3); carbon monoxide dehydrogenase (coxL-I). (**C**) CAZy enzyme Simpson diversity distributions for genomes. Simpson diversity has been transformed to the inverse form (1/(1-simpson)) for ease of viewing. Points are colored by phylum (see Fig. 3). A black star between box plots indicates a significant statistical difference (p <0.05; Wilcoxin test). Also, see Supplementary Tables 14-18.

Genomes at shallower depth were significantly enriched in C1 metabolism including: methanol oxidation (xoxF; p = 3.7e^-4^), formaldehyde oxidation (Fal1; p = 6.1e^-4^ and Fal2; p = 4.4e^-7^), methylamine to formaldehyde oxidation (mauAB; FDR = 2.8e^-9^), and formate oxidation (Frm2; p = 1.7e^-7^) (Fig. 4b and Supplementary Table 15). These organisms also had more diverse CAZy inventories compared to those at depth (Fig. 4c and Supplementary Table 16). CAZy enzymes from 29 different classes were statistically more abundant in shallow soil, and 29% are active on various forms of starch (Supplementary Table 17).

Genomes more abundant in deeper soil were enriched in both steps for complete ammonia oxidation to nitrate (amo-pmo; p = 0.04 and nxrAB; p = .005) (Fig. 4b and Supplementary Table 15). Relative to shallow soil, only three CAZy classes were statistically more abundant in genomes of depth enriched organisms (Supplementary Table 17).

We identified a further 131 KEGG functions that were statistically overrepresented in genomes that increased and 280 that decreased with depth using Random Forest-based feature selection (methods) (Supplementary Table 18). Organisms more abundant at depth are significantly enriched in PII nitrogen regulatory proteins and organisms enriched at shallower depth had significantly higher proportions of small molecule dehydrogenases active on xanthines and succinate (Supplementary Table 18). These results indicate simple carbon metabolism is more common in organisms closer to the surface and inorganic nitrogen metabolism is more common in organisms more abundant at deeper depth.

### Extended rainfall decreases depth-based differentiation in soil microbial community composition

We tested for differential abundance of organisms between samples that did and did not receive extended spring rainfall. Sample sets collected from 10-20 cm and 30-40 cm depth were analyzed separately to control for the strong phylogenetic and metabolic signal observed with depth. With extended rainfall, at 10-20 cm, 101 organisms increased and 72 organisms decreased in abundance and at 30-40 cm, 26 organisms increased and 59 decreased in abundance (Supplementary Table 5). If an organism changed in abundance at both depths, the direction of change in 95% of cases was consistent across depths. At 10-20 cm, Acidobacteria and Bacteroidetes genomes were enriched in response to extended rainfall while Chloroflexi decreased. At 30-40 cm, Thaumarchaeota and Bathyarchaeota were enriched in the group that decreased in abundance while Verrucomicrobia Bacteroidetes and RIF-CHLX increased in abundance. Thus, archaeal lineages preferentially enriched at depth become less abundant in deeper soil in response to rainfall extension with a corresponding increase in lineages associated with complex carbon degradation (Fig 4a and Supplementary Table 14).

At 10-20 cm, the genomes of organisms that increased in abundance with rainfall addition were statistically enriched in methanol dehydrogenase (xoxF; p = 7.9e^-4^) and methylamine dehydrogenase (qhpA; p = 1.2e^-5^). Organisms that decreased in abundance were slightly enriched in formaldehyde dehydrogenase (Fal1; p = 0.01).

At 30-40 cm, no function was enriched in organisms that increased after rainfall addition, however across organisms that decreased there was over representation of a variety of ammonia producing and ammonia oxidizing processes including nitrile hydrolysis (nit; p = 0.04), urea hydrolysis (URE; p = 0.01), methylamine to formaldehyde oxidation (qhpA; p = 0.02), and ammonia oxidation (amo-pmo; p = 0.001) (Fig. 4b and Supplementary Table 15).

We also assessed the CAZy diversity of organisms that changed in abundance in response to rainfall addition. At 10-20 cm we found no difference in CAZy enzyme diversity between organisms that changed in abundance with rainfall addition (Fig. 4c). However, at 30-40 cm, organisms whose abundances increased with with added rainfall had significantly higher CAZy enzyme diversity than those that decreased in abundance (Fig. 4c and Supplementary Table 16). There were specifically 24 CAZy classes more abundant in genomes that increased at 30-40 cm, the majority of which have predicted activity on pectin and hemicellulose (Supplementary Table 17). Thus, in terms of carbohydrate degradation capacity, increased rainfall pushes deeper soil towards broader complex carbohydrate degradation potential.

## Discussion

We recovered genomes for >50% of the organisms in a grassland soil, based on coverage (which is a measure of cells sampled) (Fig. 1a), and significantly expand the availability of genomes for soil organisms from poorly sampled phyla. We provide evidence that metabolic systems for processing C1 compounds were highly abundant and phylogenetically widespread, suggesting their importance in soil (Fig. 3a). Additionally, we identified unexpected organisms that engage in inorganic nitrogen turnover as well as show that the phyla and their associated functions for carbon and nitrogen metabolism are highly stratified across soil depths. It is also evident that long term climate alteration not only shifts community composition but alters the abundance of functions for important carbon and nitrogen biogeochemical cycling reactions.

Lanthanide cofactor bearing XoxF-type methanol dehydrogenases were highly prevalent, and the only methanol dehydrogenase class identified at our site. This further supports the importance of C1 compounds as a carbon currency, and implicates lanthanides as important mediators of carbon turnover in soils. Lanthanides are often sequestered into phosphate minerals^30,31^ with low biological avaliblity^32,33^. Dissolution of insoluble lanthanide phosphates likely requires strong complexation of the lanthanide ions by molecules such as siderophores^34^. In a recent report analyzing genomes from the same site it was found that Gemmatimonadetes, Rokubacteria, and Acidobacteria that harbor large numbers of XoxF sequences also exhibit extensive capacity for secondary metabolite biosynthesis^35^. Thus, we suspect a link between the prevalence of lanthanide-requiring enzymes and capacity to biosynthesize diverse small organic molecules that promote mineral dissolution.

Finding credible type-I coxL CO dehydrogenases across a variety of phyla, including highly novel Chloroflexi lineages, supports CO as an important C1 energy source in soils^36^, and expands the range of bacteria likely performing CO oxidation. However, many coxL-like sequences identified were divergent from genuine type-I coxL sequences and likely have other substrate specificity (Supplementary Fig. 8). Many molybdoprotein dehydrogenases act on small molecules like nicotinate and succinate^37,38^, which constitute large fractions of plant exudates^39^. Thus, it is probable that many of these enzymes play critical roles in plant exudate processing and turnover in soil, and represent further untracked mediators of small molecule turnover.

Complex carbohydrate degradation was primarily associated with Acidobacteria and Bacteroidetes (Fig. 3 and Supplementary Fig. 10). Gp2 Acidobacteria, which are highly abundant in some soils^40^, encode large repertoires of CAZy enzymes (Supplementary Fig. 10), and thus may represent an important and overlooked complex carbohydrate turnover sink. In contrast, the high prevalence of carbohydrate esterases we detected as well as the co-occurrence in genomes of the capacity to process acetate support the hypothesis of C1 compounds and small organic molecules being important and readily available carbon currencies for diverse organisms. As methyl and acetyl groups are common additions to a wide array of polymers^27,28^ the widespread prevalence of carbohydrate esterases may represent a strategy where readily available C1 and C2 carbon is accessed with minimal energetic investment. This observation may explain, in part, why low molecular weight carbon molecules are important currencies in this ecosystem.

The observation that most organisms with the capacity for inorganic nitrogen turnover only encode single steps of these pathways (Fig. 3b) parallels a similar finding for complex subsurface microbial communities^22^. Thus, both soils and sediments may be structured by metabolic handoffs, leading to high degrees of inter-organism cooperativity. However, only a small subset of organisms in our soils control the final steps of denitrification, and thus affect release of the climate-change relevant gasses N_2_O and NO from this system.

We found that grassland soils can be highly stratified both phylogenetically and functionally. Additionally, deeper soils were significantly enriched in organism groups that generally have low levels of genomic representation in databases. These findings have broad implications for understanding soil organic matter (SOC) turnover, as it is known that deeper strata account for a much larger fraction of SOC, with a much longer turnover time, than SOC in shallow soil^41^. The genomes reported here thus contribute significantly to understanding of bacteria and archaea that could exert critical controls on turnover rates of carbon stored in deeper soils.

The enrichment of enzymes involved in complex carbon metabolism, C1, and small molecule turnover in organisms closer to the surface (Fig. 4b-c) suggests that metabolic strategies at shallow depths are structured around plant-derived exudates and complex carbon. These data support the observation that SOM has significantly shorter residence time closer to the soil surface^41^. In contrast, most inorganic nitrogen transformation functions are more prevalent or exclusively found in organisms enriched at greater depth (Fig. 4b). Thus, deeper grassland soils could be the predominant source of N_2_O discharged to the atmosphere.

Under extended rainfall, the decrease in organisms at deeper depths performing ammonia liberation and oxidation suggests a mechanism by which climate change could directly impact subsurface nitrogen cycling and N_2_O release. These functional shifts were also joined by an increased complex carbohydrate degradation capacity at depth. The enrichment in complex carbon compound metabolism and decrease in ammonia oxidation at depth with extended spring rain suggests that deeper soils will become more favorable for carbon utilization. While the kinetics of CO_2_ release in response to these changes remains uncertain, a climate driven increase in carbohydrate degradation capacity at deeper soil depth may have important implications for the turnover of SOM.

Overall we have significantly expanded the genomic understanding of poorly characterized lineages from grassland soil environments, particularly at lower depth strata. From our work it is clear climate change also has a direct impact on organisms in this system important for trace gas cycling. Future efforts connecting this genomic information with trace gas and nutrient flux kinetics will allow a broader understanding of the contribution of grassland soils to the global biosphere.

## Acknowledgements

We thank Sue Spaulding for assistance with fieldwork and Evan Starr for helpful discussions on data analysis and figure production. Sequencing was carried out under a Community Sequencing Project at the Joint Genome Institute. Funding was provided by the Office of Science, Office of Biological and Environmental Research, of the US Department of Energy Grant DOE-SC10010566

## Data Availability

Genomic data including assemblies and raw reads will be made available under the NCBI BioProject accession number PRJNA449266. Code involved in analysis is available at the following link: https://github.com/SDmetagenomics/Angelo2018_Paper

